# The chromatin landscape of the euryarchaeon *Haloferax volcanii*

**DOI:** 10.1101/2022.07.22.501187

**Authors:** Georgi K. Marinov, S. Tansu Bagdatli, Tong Wu, Chuan He, Anshul Kundaje, William J. Greenleaf

## Abstract

Archaea, together with Bacteria, represent the two main divisions of life on Earth, with many of the defining characteristics of the more complex eukaryotes tracing their origin to evolutionary innovations first made in their archaeal ancestors. One of the most notable such features is nucleosomal chromatin, although archaeal histones and chromatin differ significantly from those of eukaryotes. Despite increased interest in archaeal histones in recent years, the properties of archaeal chromatin have been little studied using genomic tools. Here, we adapt the ATAC-seq assay to archaea and use it to map the accessible landscape of the genome of the euryarchaeote *Haloferax volcanii*. We integrate the resulting datasets with genome-wide maps of active transcription and single-stranded DNA (ssDNA) and find that while *H. volcanii* promoters exist in a preferentially accessible state, unlike most eukaryotes, modulation of transcriptional activity is not associated with changes in promoter accessibility. Applying orthogonal single-molecule footprinting methods, we quantify the absolute levels of physical protection of *H. volcanii*, and find that *Haloferax* chromatin is similarly or only slightly more accessible, in aggregate, than that of eukaryotes. We also evaluate the degree of coordination of transcription within archaeal operons and make the unexpected observation that some CRISPR arrays are associated with highly prevalent ssDNA structures. These results provide a foundation for the future functional studies of archaeal chromatin.

## Introduction

Life on earth is now understood to be divided into two deep fundamental clades – Archaea and Bacteria. Archaea were only discovered as a separate branch of the tree of life in the 1970s ^1^, yet it was noticed very early on that they share a number of common features with the more organizationally complex eukaryotes, especially in the organization of their information processing cellular machinery. Based on these similarities it was suggested that eukaryotes evolved from archaea ^2,3^, a view strengthened in the phylogenomic era ^4^, and eventually solidified with the discovery of archaeal lineages such as the Lokiarchaeota ^5^. Thus, we now know that many of the complex cellular features that characterize eukaryotes trace their origins to their archaeal ancestry ^6,7^.

One of the most notable such features is nucleosomal chromatin. Nearly all eukaryotic genomes are packaged by nucleosomes, consisting of two tetramers of the four core histones H2A, H2B, H3 and H4, wrapping around ∼147 bp of DNA. These proteins are, with very rare exceptions ^8,9^, the most evolutionarily conserved among eukaryotes ^10^, in large part because aside from their packaging function they are also subject to a large number of precisely regulated posttranslational modifications (PTMs) at key residues ^11^, through which they play a pivotal role in all aspects of chromatin biology (transcription and its regulation, DNA replication, DNA repairs, mitosis, and others).

As early as the 1980s it was noticed that some archaea possess proteins and structures similar to eukaryotic histones and nucleosomes ^12,13^. We now know that most archaea have nucleosomal chromatin and histones ^14–16^, and that these histones are ancestral to the eukaryotic histones. This is not true for all archaea as some lineages use other proteins such as Alba/Sac10b, Sul7d, Cren7, and CC1^17,18^ to package their genomes, but the most often observed state is to have histones.

However, archaeal histones differ substantially from those in eukaryotes – while they share the core histone fold domain, they usually do not have the unstructured tails of H2A/H2B/H3/H4 that are the main sites of key PTMs. Archaeal histones also do not form octameric nucleosomes; instead, only one or a very small number of histone genes are found in archaeal genomes, and the structures they form are very different from those of eukaryotes. The diversity of histone sequences across the whole archaeal phylogeny is very large and still largely unexplored experimentally, but the available structural ^19^, biochemical and modeling work suggests that in at least some species histones can form so called “hypernucleosomes” or “archaeasomes”, consisting of a protein core of individual histones stacked next to each other, around which DNA is wrapped ^15,20^, ranging from 60 to 500 bp ^21^. It has also been proposed that archaeal histones exhibit an inherently dynamic association with DNA, resulting in so called chromatin ‘slinkies’ that can easily slide along DNA ^21^, in contrast to the much more stable association of nucleosomes with DNA in eukaryotes.

Despite the relevance of archaeal chromatin to understanding the deep evolution of chromatin organization, up until now the structure of archaeal chromatin has received direct experimental investigation using modern genomic tools, with the exception of early MNase-seq studies nearly a decade ago that mapped nucleosomal positioning in the euryarchaeotes *Haloferax volcanii* ^22^ and *Methanothermobacter thermautotrophicus* and *Thermococcus kodakarensis* ^23^. Furthermore, very little is known about the relationship between chromatin structure and the regulation of gene expression in these organisms.

The picture has become even more complicated in recent years by the suggestion that in some archaea, in particular haloarchaea, the presence of histone proteins does not necessarily mean that chromatin is packaged by them ^24–29^. This tentative conclusion is based on the low abundance of the HpyA protein in *Haloferax* proteomics datasets ^28,29^ and on data suggesting that it is not necessarily involved in genome packaging but has functions analogous to a transcription factor ^26^. Thus the precise nature of chromatin structure in these archaea remains an open question.

To fill these gaps in our understanding of the organization of archaeal chromatin, we mapped chromatin accessibility and active transcription in *Haloferax volcanii* using a combination of bulk and single-molecule techniques such as ATAC-seq ^30^ (Assay for Transposase-Accessible Chromatin using sequencing), NOMe-seq/dSMF ^31^ (Nucleosome Occupancy and Methylome sequencing/dual Single-Molecule footprinting) and KAS-seq ^32^ (Kethoxal-assisted single-stranded DNA sequencing). We find that chromatin in *H. volcanii* exhibits similar features to that of eukaryotes on a broad level, with preferentially accessible promoters regions. However, unlike in eukaryotes, chromatin accessibility at promoters does not relate to transcriptional activity. Using single-molecule footprinting we estimate absolute protein occupancy levels over the *H. volcanii* to be comparable to, or possibly slightly lower than those in eukaryotes. However, unlike what is seen in most eukaryotes, we do not observe stably positioned nucleosome protection footprints, but rather only statistically elevated accessibility around promoters. We also revisit the question about the degree of coordination of transcriptional activity and chromatin accessibility within *Haloferax* operons, and make the unexpected discovery that some CRISPR arrays are associated with very strong ssDNA signatures.

## Results

### ATAC-seq reveals the open chromatin landscape of *H. volcanii*

To study chromatin accessibility in archaea we adapted the ATAC-seq assay ^30^ to the *Haloferax volcanii* archaeon. *H. volcanii* is a halophile with a strong preference for very high salt concentrations in the growth medium (see the Methods section), which grows optimally at 42 °C ^33,34^, and is a widely used archaeal model system.

The principle behind ATAC-seq is the very strong preference of the Tn5 transposase ^30^ for insertion into accessible DNA as opposed to tagmentation of protected (by nucleosomes, transcription factors, or other proteins) DNA. Tn5 insertion then tags accessible sites with landing sites for PCR primers, allowing for highly efficient amplification of open chromatin regions in the genome.

After extensive testing of a variety of different experimental protocols (fixation conditions and input cell numbers), we arrived at the following modifications of the standard ATAC protocol. First, because archaea are not eukaryotes and do not have a nucleus, we omitted the cell lysis and nuclei isolation step that is a standard feature of eukaryotic ATAC-seq protocols, such as the now standard omniATAC^35^. Second, and most important, we reasoned that if the previously reported dynamic repositioning of archaeal nucleosomes along DNA occurs in *H. volcanii*, optimal results might be obtained by introducing a crosslinking step into the standard ATAC protocol, which would “freeze” any nucleosomes and other structural proteins in place and not allow transposition into DNA that might change from protected to accessible during the duration of the transposition reaction. Indeed, comparing the TSS (transcription start site) enrichment generated without fixation and with light (0.1% formaldehyde) and strong (1% formaldehyde) fixation showed that strong fixation produces stronger TSS enrichment (Figure 1A). We then compared the *H. volcanii* ATAC-seq TSS metaprofile with that from the previously published MNase-seq dataset and observed the expected inverse relationship (Figure 1B). *H. volcanii* TSSs exhibit elevated accessibility in the 0 to −400 bp upstream region, decreasing away from the TSS, and displaying a weak periodicity of ∼60-70 bp.

**Figure 1:**
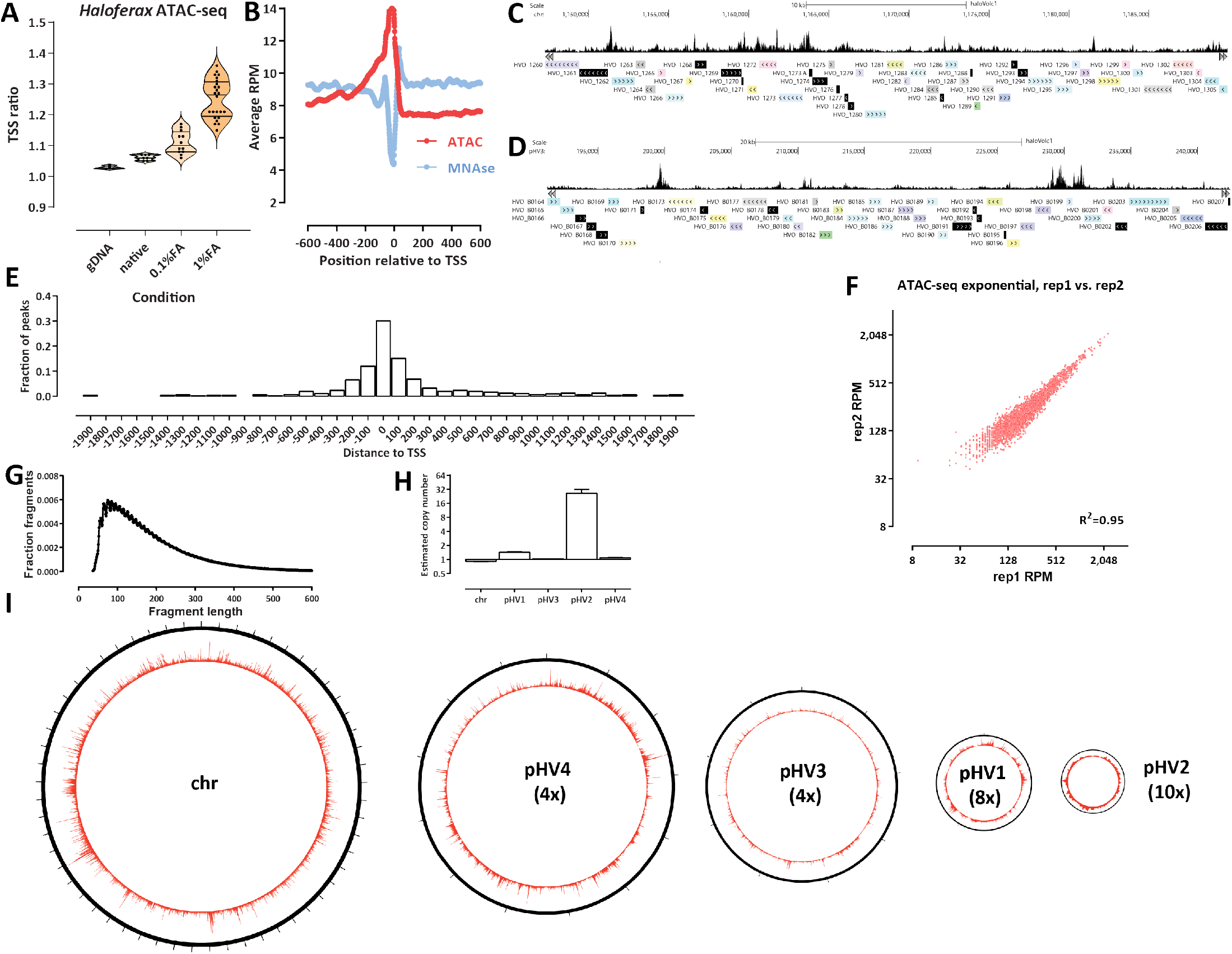
Archaeal ATAC-seq and the open chromatin landscape of *H. volcanii*. (A and B) Adaptation and optimization of the ATAC-seq assay to the archaeal context. (A) Distribution of TSS ratio scores (see Methods for details) for native, 0.1%- and 1%-formaldehyde ATAC-seq libraries. (C and D) Representative browser snapshots of ATAC-seq profiles along the *H. volcanii* genome. (E) Distribution of MACS2 ATAC-seq peaks relative to TSSs. (F) High reproducibility of *H. volcanii* chromatin accessibility measurements using ATAC-seq. Shown is the between-replicate correlation over TSSs in RPM (Read Per Million) units. (G) Fragment length distribution in *H. volcanii* ATAC-seq datasets. (H) Estimated relative copy number of *H. volcanii* chromosomes. Genomic DNA was tagmented and amplified (*n* = 4) and normalized read coverage was estimated for each chromosome/plasmid. The average ratios are shown. (I) Global ATAC-seq profile over each of the five *H. volcanii* chromosomes. The number in brackets corresponds to the magnification of the true proportional size of plasmids relative to the main chromosome.

Genome browser visualization of ATAC-seq profiles (Figure 1C-D) revealed an accessibility landscape largely reminiscent of that in eukaryotes with compact genomes such as the budding yeast *Saccharomyces cerevisiae* ^36^, with accessibility concentrated at TSSs. Peak calling using MACS2^37^ generalized that observation (Figure 1E) – nearly all ATAC-seq peaks are located within 200 bp of an annotated TSS.

ATAC-seq measurements in *H. volcanii* are also highly reproducible between experimental replicates (Figure 1F).

The *H. volcanii* ATAC-seq fragment length distribution is unimodal, peaking at 90-100 bp, and does not show the eukaryotic mono-, di- and tri-nucleosomal signature (Figure 1G).

The *H. volcanii* genome consists of multiple replicons ^38^, with a main chromosome (“chr”) and four plasmids of very different size – pHV4, pHV3, pHV1 and pHV2 (in order of decreasing size), which together comprise ∼30% of the total genome. To properly interpret sequencing data (which is typically normalized to total read coverage), we determined the relative copy number distribution of these replicons using tagmented naked genomic DNA control samples (Figure 1H). The smallest plasmid – pHV2 – appears to exist in ∼26 copies for each main chromosome, pHV1 is found at ∼1.4 copies for each main chromosome, while the two large plasmids – pHV4 and pHV3 – exist in a 1:1 ratio to the main chromosome.

As previous studies of chromatin openness in bacteria have reported the existence of very large domains of lower and higher accessibility ^39^, we wondered whether the same is observed in *Haloferax*. We do not observe such domains in our datasets (Figure 1I). Recently, an ATAC-seq dataset was reported from the crenarchaeote *Sulfolobus islandicus* ^40^, which lacks histones, and instead packages its genome mainly through Alba/Sac10b proteins ^41,42^. In that species, large domains similar to those in bacteria were reported, suggesting that such large-scale domains of elevated chromatin accessibility might be a feature associated with the lack of nucleosomal chromatin in prokaryotes, while archaea that contain histone genes, such as *H. volcanii*, exhibit eukaryote-like organization. We also reexamined the *Sulfolobus islandicus* dataset and found that it displays a much more modest TSS enrichment than that seen in *H. volcanii*, which is also more narrowly concentrated around the TSS position (Supplementary Figure 1).

Finally, we observed an anti-correlation between ATAC-seq signal and genomic GC content (Supplementary Figure 2). *Haloferax volcanii* exhibits a rather high average GC content of 65% ^38^, decreasing to 58% in intergenic regions, and areas with even higher GC content (*>*65%) show markedly lower ATAC-seq signal. This observation is corroborated by the available external MNase-seq dataset, which shows positive correlation with GC content (i.e. the inverse of ATAC-seq, as expected) and naked DNA controls (which show no correlation with GC content, indicating that PCR biases during sequencing library preparation are not the reason for the observed patterns).

### Absolute DNA occupancy/protection levels in *H. volcanii*

While ATAC-seq is immensely helpful for identifying the location of accessible regions in the genomes and measuring their relative accessibility, it is a bulk method that does not provide information about the absolute levels of protection/accessibility in the genome. Instead, absolute accessibility must be measured either restriction digestion-based or enzymatic labeling single-molecule methods. To quantify absolute occupancy/protection levels in the *H. volcanii* genome, we applied NOMe-seq ^43^ and dSMF ^31^ to *Haloferax* chromatin. These methods rely on the preferential methylation of accessible cytosine nucleotides (5mC) by a recombinant methyltransferase that modifies specifically in GpC contexts (NOMe-seq) or a combination of methyltransferases that label both GpC and CpG (dSMF).

However, these methods are potentially confounded by the presence of endogenous methylation in either context. Fortunately, in the case of *H. volcanii* endogenous DNA modifications have been previously studied using PacBio single molecule sequencing, and no CpG and GpC modifications were found. Instead, only two restriction modification system-associated modifications in different contexts, specifically, 4-methylcytosine in a C(m4)TAG context and N6-methyladenine in a GCA(m6)BN6VTGC context were identified ^44^.

Figures 2A-C show the metaprofiles of average methylation around *H. volcanii* TSSs for NOMe-seq and dSMF datasets generated from exponentially growing and stationary cultures (post log-phase in the growth curve). We observe baseline absolute protection levels around 84-85% in the exponentially growing cells and ∼ 89% in stationary cells. For comparison, analogous studies in eukaryotes, such as the budding yeast *S. cerevisiae* ^36,45^, have shown absolute protection levels around 90% (±5%). Thus, *Haloferax* chromatin exhibits broadly similar, though perhaps somewhat lower levels of protection than what is observed in conventional eukaryotes.

**Figure 2:**
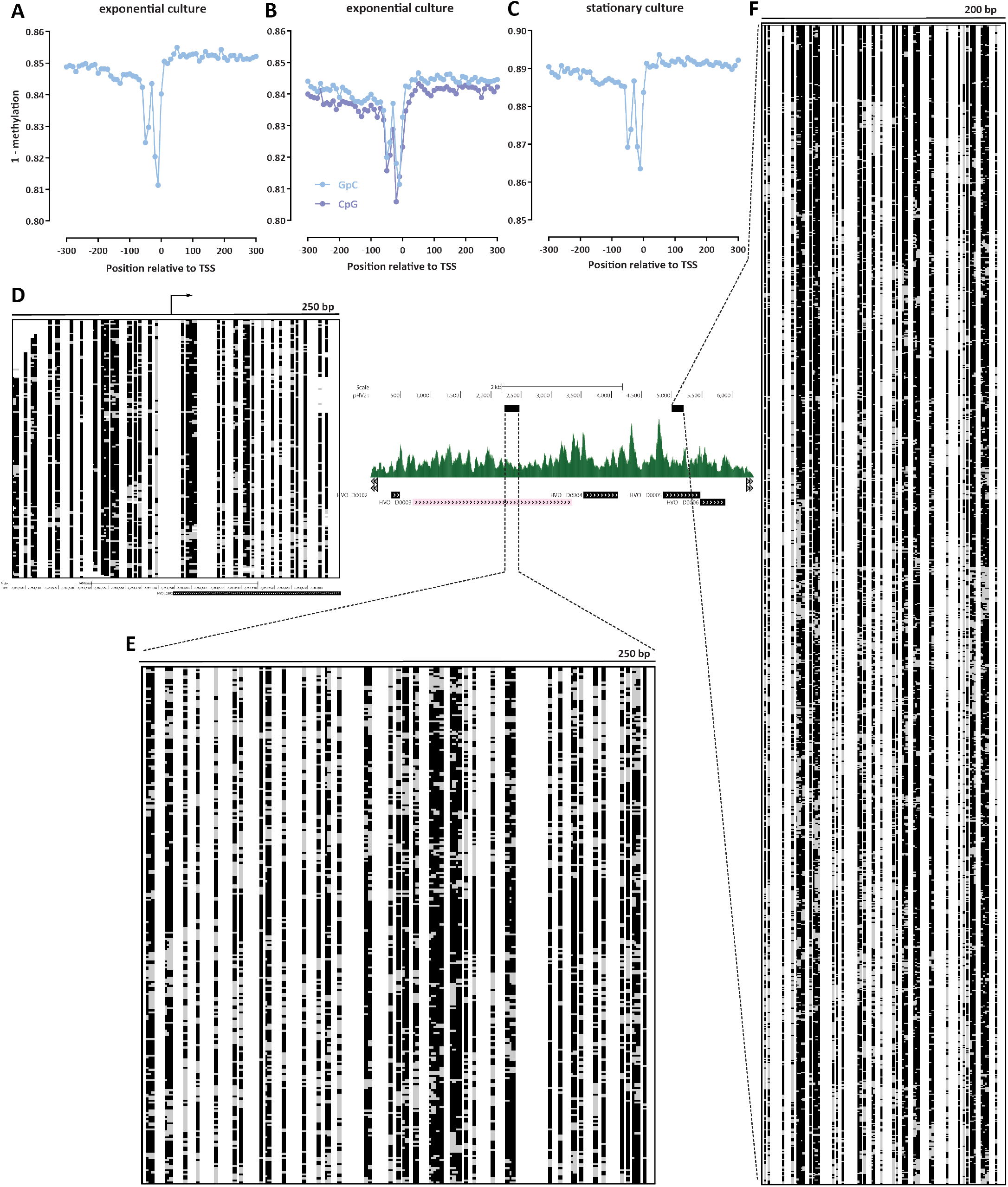
Absolute DNA occupancy/protection levels in *H. volcanii*. (A-C) TSS metaprofiles in different conditions (two replicates of an exponentially dividing culture, and a stationary culture). (D) Single-molecule map (250-bp) around a main-chromosome TSS. Black indicates unmethylated and therefore protected sites, gray indicates methylated and thus accessible sites. (E and F) Single-molecule maps over the pHV2 plasmid: 250 bp-window map (E) and very high coverage (≥1,200 single molecules) 200 bp-window map (F).

The base-pair resolved nature of these single-molecule methods enabled an observation of a feature not readily apparent in ATAC-seq and MNase-seq datasets – a protection footprint immediately upstream of the TSS. We also observe this feature as a protection footprint in a few percent of single molecules at individual promoters (Figure 2D). At present we are not able to confidently identify its functional association – its width is likely too small for it to be a positioned −1 nucleosome, and we hypothesize it may correspond to one of the complexes involved in the archaeal transcriptional cycle, analogous to how similar protection footprints associated with the RNA polymerase and the preinitiation complex (PIC) in eukaryotes have been observed in dSMF datasets ^31^.

On the other hand, unlike this unique protection footprint, we do not observe strongly positioned individual nucleosome-like features along the *Haloferax* genome, such as those seen in conventional eukaryotes (Figure 2D). However, our NOMe-seq and dSMF datasets were sequenced at an effective depth of ∼ 100 × for fragments of width 200 bp. To determine if higher sequencing depth would clearly reveal such putative positioned nucleosomes (or other proteins), we turned to the pHV2 plasmid, which, as previously discussed, exists in high copy numbers in *H. volcanii* cells. Over the pHV2 plasmid we obtained ∼200× coverage for fragments of length 250 bp (Figure 2E) and ∼1,200× coverage for fragments of length 200 bp (Figure 2F). These high-depth maps also reveal considerable heterogeneity of footprints and accessible sites. These observations are consistent with previous proposals for highly dynamic association of archaeal nucleosomes with DNA, but also with packaging by other, non-histone proteins that is similarly positionally unstable.

### The ssDNA and active transcription landscape in the *H. volcanii* genome

We then turned our attention to the landscape of active transcription in *H. volcanii*. To this end, we used KAS-seq ^32^, which measures with high specificity the presence of single-stranded DNA in the genome, by labeling unpaired guanine nucleotides with N_3_-kethoxal, using click-chemistry to add a biotin moiety, then enriching these fragments via a streptavidin pulldown. Most ssDNA is usually found within the transcriptional bubbles associated with RNA polymerase molecules engaged with DNA ^32^. KAS-seq provides several advantages in the *H. volcanii* context. First, due to the absence of straightforward methods to deplete *H. volcanii* ribosomal RNA (rRNA) from RNA sequencing libraries analogous to polyA-selection in eukaryotes ^47,48^, it enables measurement of transcriptional activity at much lower cost than deep RNA-seq experiments. Second, and most important, it measures actively engaged polymerase molecules, and thus unlike the steady-state transcript levels that conventional RNA-seq quantifies, it provides a means to quantify active transcriptional activity. Third, it also identifies other ssDNA structures, such as those resulting from paused polymerase molecules, G-quadruplexes, and others.

We carried out a time course analysis of *Haloferax* growth and applied both KAS-seq and ATAC-seq during the “exponential” log-phase of growth, and on the “stationary” post-log phase stage, as well as on “standing” cultures, which had been left at room temperature for ∼1 week. We also carried out KAS-seq on exponentially growing cells that were then incubated at different temperatures – the typical growing temperature of 42 °C, 37 °C, 23 °C and a cold shock at 4 °C for 4 hours.

At a global level, we observe uniform levels of KAS signal along the length of *H. volcanii* chromosomes (Figure 3A), with sharp localized peaks. KAS-seq measurements in *H. volcanii* are highly reproducible between experimental replicates (Figure 3B). Locally, at the level of individual genes, we observe a combination of high-signal peaks at the promoters of some genes and elevated KAS-seq signal along gene bodies (Figure 3C-D). This feature is also in KAS-seq metaprofiles over all genes (Figure 3E), suggesting that in *H. volcanii* RNA polymerases spend substantial amount of time associated with the TSS, perhaps in a paused state analogous to that observed in metazoans ^49^ or the archaeon *Sulfolobus solfataricus* ^50^.

**Figure 3:**
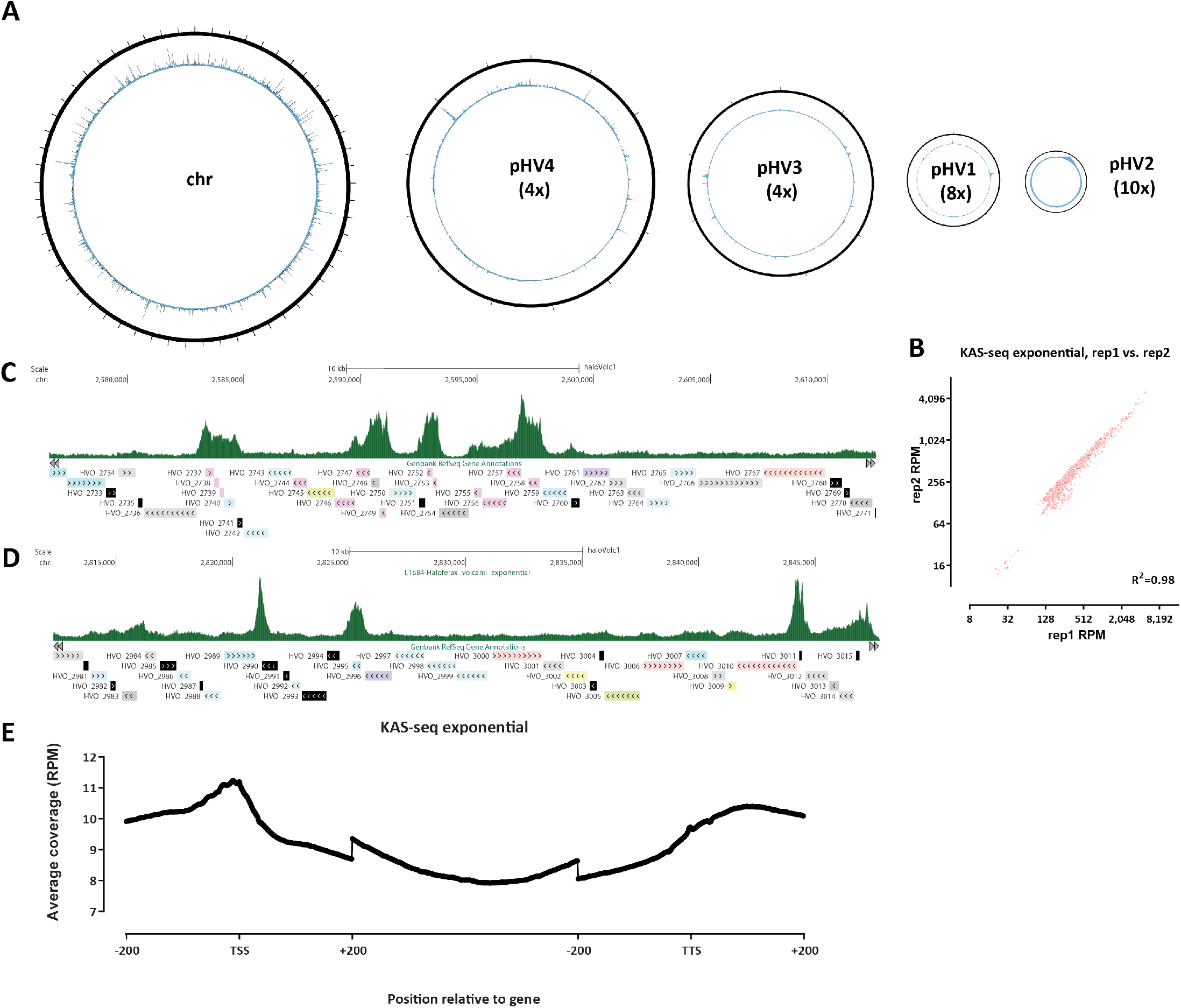
The ssDNA and active transcription landscape in the *H. volcanii* genome as measured by KAS-seq. (A) Global KAS-seq profiles over each of the five *H. volcanii* chromosomes in an exponential culture. (B) High reproducibility of active transcription measurements using KAS-seq. (C-D) Representative browser snapshots of KAS-seq profiles along the *H. volcanii* genome. (E) KAS-seq metaprofile along *H. volcanii* gene bodies.

### Strong, culture condition-dependen ssDNA signals are associated with some *H. volcanii* CRISPR arrays

In the process of optimization of the KAS-seq assay in *Haloferax*, we carried out KAS-seq on a *H. volcanii* culture that had been left standing at room temperature for 3∼ months. These data revealed that CRISPR arrays can become highly enriched for KAS-seq signals in these viable, yet dormant, cultures. CRISPR (clustered regularly interspaced short palindromic repeats) arrays are a key element in the defense systems against foreign genetic material of many prokaryotes and consist of multiple identical repeats interspersed with non-repetitive sequences that target foreign plasmids and phages, together with a set of Cas genes. *H. volcanii* is one of the prokaryotic systems where these elements were first originally observed ^51–53^.

In standing *Haloferax* cultures, transcriptional activity is likely suppressed, as the cells enter a dormant state. Consistent with a transcriptionally inactive state, we observe largely flat, low levels of KAS-seq signal from this standing culture KAS-seq dataset (Figure 4A). However, we also observed a single large, sharp peak of KAS-seq signal on the pHV4 plasmid. This peak resides between the second (as numbered in the available genome annotation) CRISPR array in the *H. volcanii* genome ^54^ and its associated Cas6 gene (Figure 4B); a previously reported small RNA (sRNA) – s479^55^ is also located in between Cas6 and the CRISPR array, but our observed KAS-seq peak is not associated with this putative promoter, but is instead situated downstream of the array. While this KAS-seq signal peak is also found in all other conditions we assessed, but it stands out in the long-term standing culture due to the absence of other peaks that result from the active transcription of other genes.

**Figure 4:**
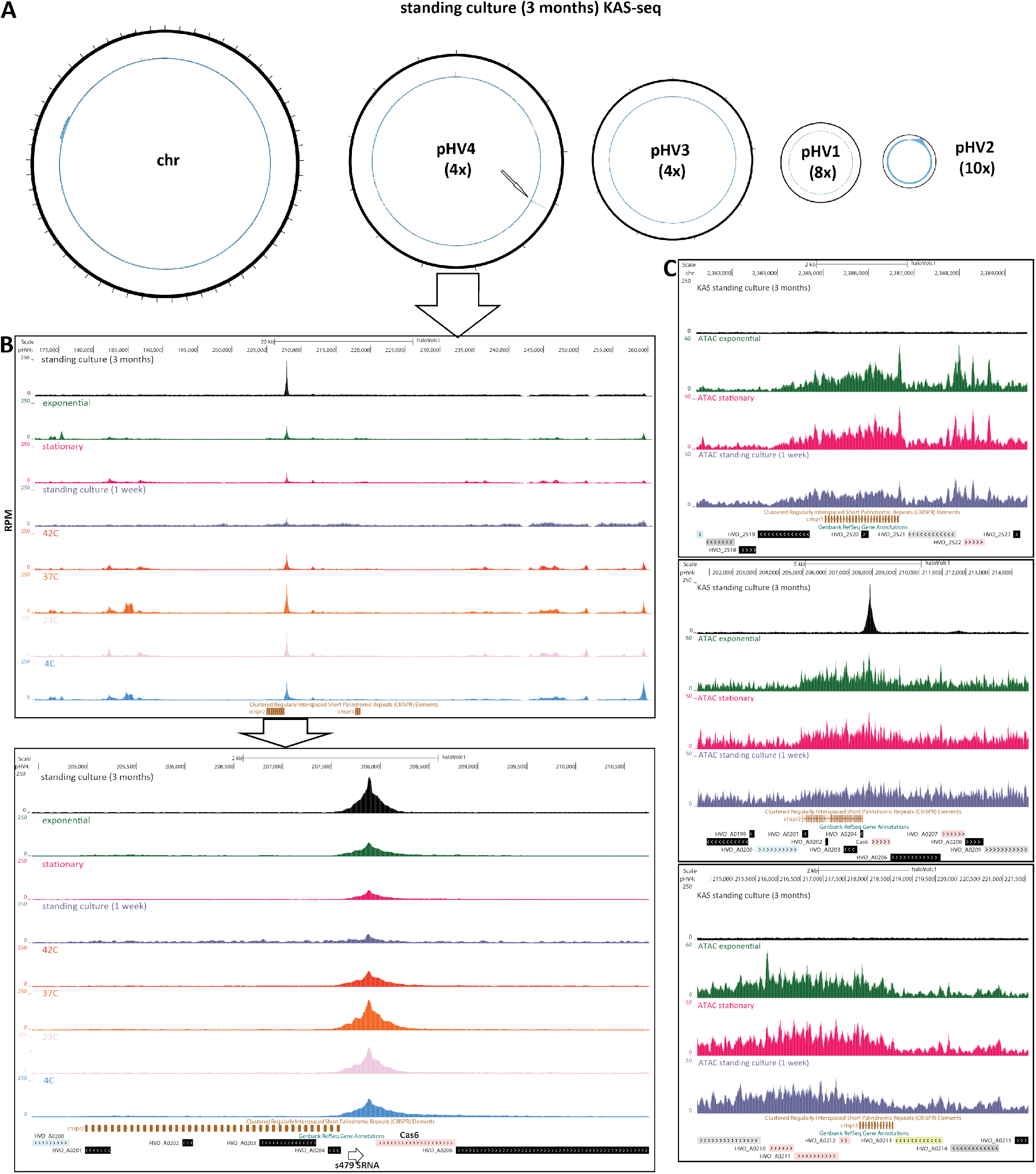
Abundant ssDNA structures associated with some *H. volcanii* CRISPR arrays in specific conditions. (A) Global KAS-seq profiles over each of the five *H. volcanii* chromosomes in a long-standing culture (∼3 months) reveals an extremely strong ssDNA peak associated with one of the CRISPR arrays on the pHV4 plasmid. (B) KAS-seq signal levels around pHV4 plasmid CRISPR arrays in different condition. (C) KAS-seq and ATAC-seq levels around all three *H. volcanii* CRISPR arrays in different conditions.

Curiously, only the second CRISPR array in *H. volcanii* displays this strong ssDNA structure, while the other two do not (this is true across all assayed conditions). However, all three arrays show elevated chromatin accessibility in ATAC-seq datasets – accessibility signal that is not focused on the beginning of the array but covers its whole length (Figure 4C). Possible interpretations of these observations are discussed below.

### Coordination between chromatin accessibility and transcriptional activity within *H. volcanii* operons

Being prokaryotes, archaea often have genes organized into operons ^56^, with multiple genes transcribed as a single unit, which are therefore expected to share a common promoter and to exhibit similar levels of active transcription. However, transcription of these operons is still little studied using modern genomic tools. To address this gap, we used our KAS-seq and ATAC-seq data, which provide information about the chromatin accessibility and active transcription in different conditions, to investigate the extent of coordination between the transcriptional activity of different units in operons in the *H. volcanii* genome.

We first inspected the two *H. volcanii* operons that include rRNA genes ^57^. Figure 5A shows KAS-seq and ATAC-seq profiles along rRNA genes in the different conditions we assayed. We observe a largely uniform KAS-seq profile in exponentially growing cells, and generally elevated chromatin accessibility (which might be associated with very active transcription); a more non-uniform pattern is seen in cold-shocked cells kept at 4 °C, where lower KAS-seq levels are seen over the large subunit (LSU) rRNA relative to the small subunit (SSU), with various intermediate states in other conditions.

**Figure 5:**
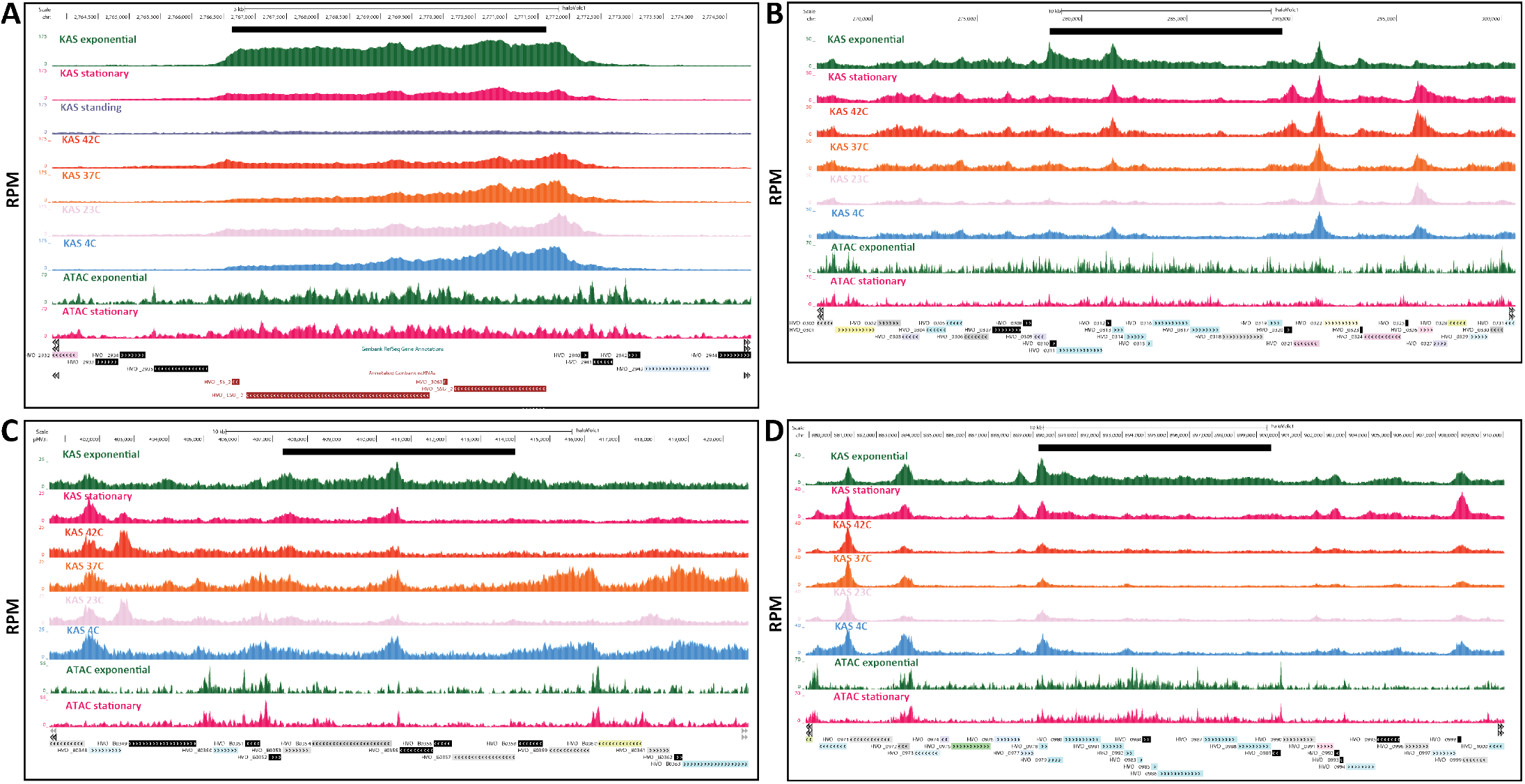
Coordination between chromatin accessibility and transcriptional activity within *H. volcanii* operons. Black bar shows the operon boundaries. (A) Ribosomal RNA operon. Note that the tracks shown here were generated by including multimapping reads (see Methods for details). (B) A-type ATP synthase subunits A, A, B, C, D, E, F, I, K, and H. (C) RNA polymerase II subunits. (D) NADH dehydrogenase-like complex subunits A, B, CD, H, I, J1, J2, K, L, M, N.

More interesting patterns are seen in operons comprised of protein coding genes. Figure 5B shows a gene array consisting of A-type ATP synthase subunits, for which distinct KAS-seq peaks are seen at the beginning of the operon as well as in between genes in the middle of the operon. Furthermore, KAS-seq levels are not uniform over the gene bodies of all genes.

We examined multiple other operons (Figure 5C-D) and Supplementary Figure 3, which reveal a very diverse picture of the extent of coordination between the transcriptional activity over individual genes within an operon – internal operon peaks are observed for multiple operons, while there are also other operons where KAS-seq signal is more uniform.

In some cases (e.g. Figure 5C), these internal KAS-seq peaks are also associated with matched ATAC-seq peaks. Thus, one interpretation of these observations is that not all these operons are true operons (even though they consist of functionally related genes), but instead independent initiation and regulation of transcription from internal TSSs may be occurring. This interpretation is particularly supported in the cases where ATAC-seq peaks are seen at the beginning of genes, and where the baseline gene-body KAS-seq signal differs greatly between different sections of the operon, and is also consistent with previous observations of internal promoters inside archaeal operons based on transcript 5’-end mapping ^47,58,59^. On the other hand, internal KAS-seq peaks might also arise from very strong and immediate coupling between transcription and translation, i.e. if the process of initiation of translation at internal positions in the operon somehow leads to the polymerase pausing at certain sites.

### Chromatin accessibility does not correlate with transcriptional activity in *H. volcanii*

In eukaryotes, the regulation of chromatin accessibility at regulatory elements (promoters and enhancers) is key to gene regulation, as nucleosomal chromatin is generally refractive to occupancy by regulatory proteins and to active transcription ^60^, and while perfect correlation between accessibility levels at promoters and gene expression is rare, open chromatin states are generally associated with increased transcriptional activity.

In contrast, the relationship between chromatin accessibility and transcriptional activity in archaea has not been systematically studied as chromatin accessibility has not been mapped globally, across conditions, and in conjunction with global measurements of active transcription.

We first identified differentially accessible promoter regions between the different conditions we studied (Figure 6A). In contrast to an a priori expectation that changes in gene expression would be associated with shifts in chromatin accessibility levels around promoter regions, we did not find strong changes between exponentially growing and stationary cells (Figure 6A). We observed large apparent differences in ATAC-seq signal in each of those two conditions and standing cultures (Supplementary Figure 4), but in those comparisons the profiles are highly skewed towards increased higher accessibility in the actively growing and stationary cells instead of showing the typical more symmetric changes (i.e. accessibility increasing and decreasing in both directions) between two conditions;

**Figure 6:**
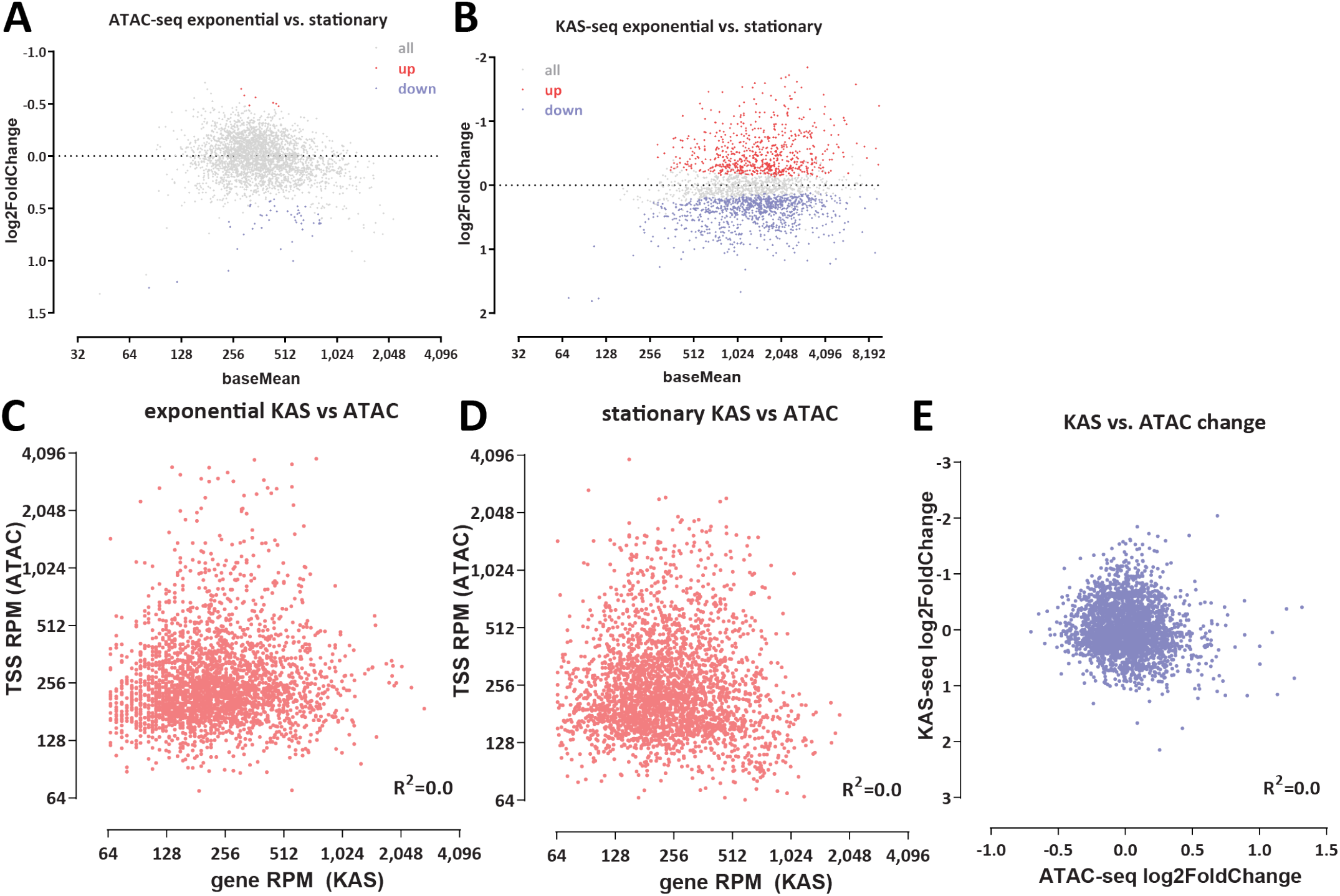
Chromatin accessibility does not correlate with transcriptional activity in *H. volcanii*. (A) Differential chromatin accessibility between exponential and stationary conditions; (B) Differential KAS-seq levels between exponential and stationary conditions; (C-D) Lack of correlation between KAS and ATAC signals in exponential and stationary conditions; (E) Lack of correlation between changes in chromatin accessibility and changes in transcriptional activity.

We believe this pattern is due to the dormant state of standing cultures in which we do not observe as strong ATAC-seq peaks as are observed in actively growing cells.

In stark contrast to the lack accessibility changes between non-dormant conditions, we find a large number of genes that display strong differential KAS-seq signal over their gene bodies (Figure 6B). Thus while we observe no major changes in chromatin accessibility across these two conditions, we do note large-scale changes in RNA polymerase occupancy (as inferred by KAS-seq).

We then quantified the degree of correlation between KAS and ATAC signals (Figure 6C-D) and found no significant correlation. We also found no correlation between the level of changes in chromatin accessibility and changes in RNA polymerase association with DNA (as inferred by KAS-seq) between exponential and stationary cells (Figure 6E).

These global observations are supported by the study of individual loci. Figure 7A shows ATAC-seq levels and KAS-seq levels over all genes across different conditions, clearly indicating that these two signals are decoupled from on another. Figure 7B and C depict other examples of genes for which transcriptional activity shifts between conditions yet ATAC-seq profiles are largely identical.

**Figure 7:**
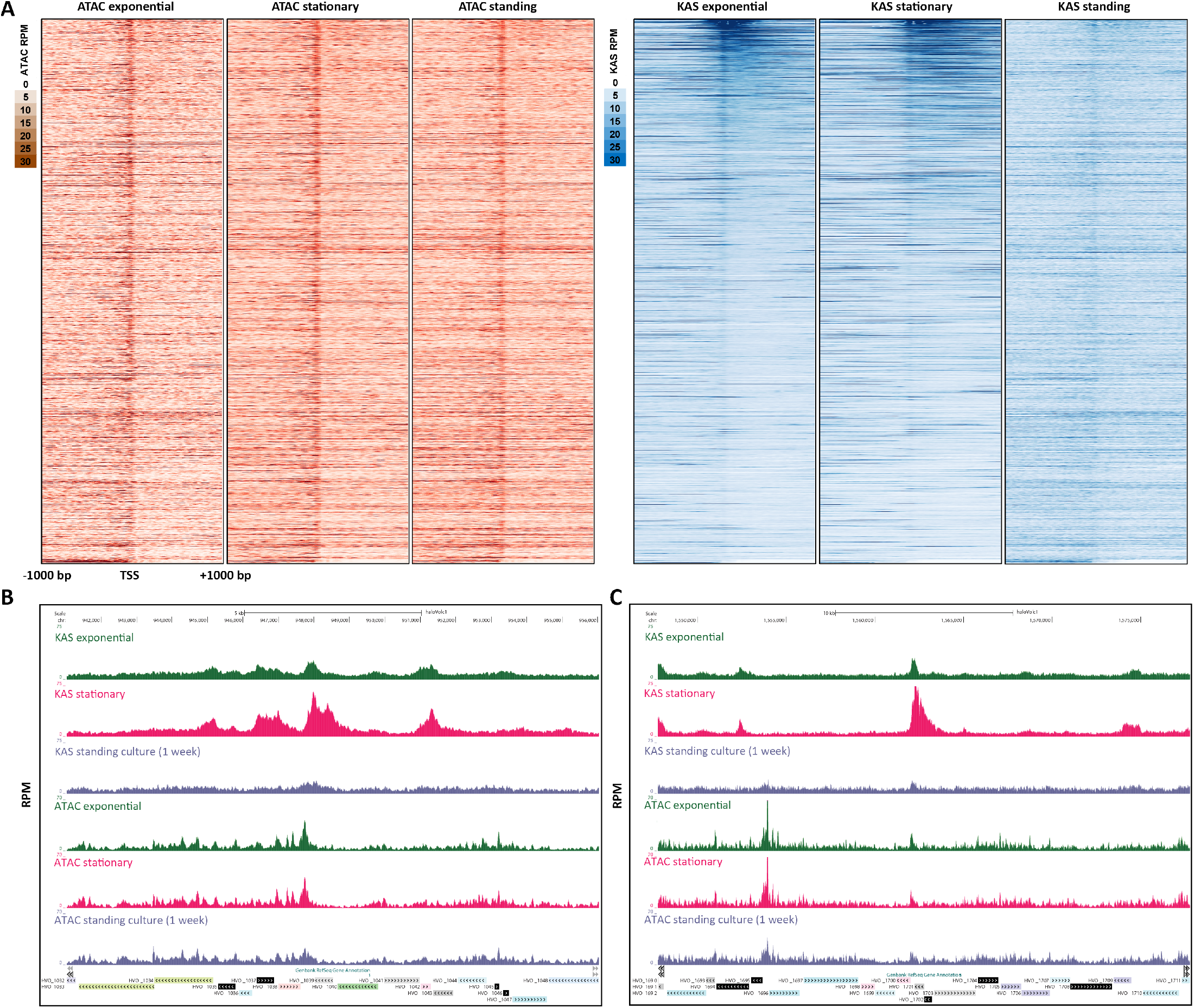
Chromatin accessibility does not correlate with transcriptional activity in *H. volcanii*. (A) Genome-wide heatmaps of ATAC-seq and KAS-seq signals around *H. volcanii* TSSs, sorted by KAS-seq levels in the exponential condition. (B-C) Representative snapshots of genes with significantly altered transcriptional activity between the exponential and standing condition, but no corresponding changes in chromatin accessibility.

We thus conclude that based on the currently available data, the modulation of chromatin accessibility does not appear to be a major determinant/correlate of transcriptional activity in *Haloferax* archaea.

## Discussion

In this study, we adapted and applied methods for global profiling of chromatin accessibility and ssDNA in the euryarchaeote *Haloferax volcanii*, revealing the chromatin architecture of this representative of the haloarchaea. We identified several convergent and divergent characteristics with respect to those of conventional eukaryotes properties.

The *H. volcanii* genome displays a chromatin organization very similar to that of eukaryotes with compact genomes such as budding yeast – accessibility peaks are almost exclusively found very close to promoters. Absolute accessibility/protection levels are similar, perhaps slightly lower than those in budding yeast, with a baseline protection level of 85-90%. On the other hand, we do not observe the strongly positioned nucleosome-like features such as those typical in eukaryotes in *Haloferax*, but instead observe a more heterogeneous picture consistent with the previously reported dynamic association of archaeasomes with DNA.

In the light of recent reports casting doubt on the nucleosomal packaging of haloarchaeal genomes ^24–26^ these observations are puzzling. Once again, the picture of chromatin accessibility revealed by ATAC-seq in *Haloferax* is nearly identical to that of conventional eukaryotes with nucleosomal chromatin and densely packed genomes. At an arbitrary locus, one would note very few general differences between between a *Haloferax* ATAC-seq signal track and a yeast ATAC-seq signal track. Both exhibit strong localized peaks at promoters, and similar levels of absolute protection/occupancy. No protein other than histones is known to confer such properties, so it is very tempting to conclude that *Haloferax* histones are responsible for the observed accessibility patterns, but this conclusion would be discordant with the most recent reports on the subject. Experiments involving *in vitro* reconstitution on DNA combined with heterologous expression in yeast of the *Haloferax* histone protein and its other putative chromatin proteins might shed light onto this question in the future.

We also note, that unlike what has been reported about bacteria and archaea without histones the *H. volcanii* genome does not exhibit large-scale domains of diminished and elevated accessibility.

In contrast to the norm in eukaryotes, accessibility at promoters does not correlate with transcriptional activity. This puzzling observation will require further functional dissecttion in future work. The idea that chromatin accessibility is not necessary for transcription in archaea is supported by previous observations that transcription by the archaeal RNA Polymerase is slowed, but not blocked by archaeal nucleosomes ^61^. However, the molecular determinants of the observed accessibility remain unclear. Nearly all promoters in *H. volcanii* show some level of accessibility (Figure 7A), but its levels differ greatly between individual genes. How these differential states are specified, and whether they might in fact change in conditions that we have not assayed remains to be determined. While MNase-seq studies *Methanothermobacter thermautotrophicus* and *Thermococcus kodakarensis* ^23^ suggested that nucleosome positioning in those organisms is significantly influenced by DNA sequence, but no such strong association was reported for *Haloferax volcanii* ^22^. In the currently available datasets we observe anti-correlation between chromatinization and high genomic GC content, but whether this is the primary determinant of nucleosomal or other protein occupancy, and whether this correlation can account for the large differences in promoter accessibility combined with a general absence of such strong peaks elsewhere in the genome observed in *Haloferax* remains uncertain. Generating chromatin accessibility and nucleosome positioning data across other archaeal clades and also in multiple closely related species will allow us to generalize these observations and to train fully powered models that relate sequence to chromatin accessibility, potentially identifying such determinants.

We also made the observation that operons in *Haloferax* display non-uniform levels of single-stranded DNA signal consistent with transcriptional activity, and may in fact consist of multiple distinct transcriptional units. The phenomenon of independent transcription of operon genes has been suggested by some previous studies in *Haloferax* and *Sulfolobus* ^62–64^, but this is again an observation and a hypothesis that needs to be generalized to and tested in not only more archaeal species, but also in bacteria, where the application of KAS-seq to the study of transcriptional activity may also result in unanticipated findings. In *Haloferax* we were only able to examine several dozen of unambiguous operons (e.g. unidirectional arrays of functionally related genes).

Bacteria are also highly relevant to the other surprising observation we made – the strong ssDNA structure present at the second *Haloferax* CRISPR array, especially in dormant cells that are otherwise mostly transcriptionally silent, but not at the other two CRISPR arrays. The second CRISPR array is uniquely associated with the Cas6 gene. Cas6 is the endoribonuclease that generates guide RNAs ^54,65^, thus one possible explanation for the strong ss-DNA peak between the CRISPR array and Cas6 is that it represents paused RNA polymerase at the Cas6 promoter. If this explanation is correct, the fact that Cas6 is the only gene in the genome with this property in dormant cells is remarkable and perhaps points to the importance of retaining the ability to process CRISPR transcripts even in a dormant cellular state. Alternatively, the ssDNA structure might be related to the transcription of the CRISPR array itself; however, such an explanation does not speak to the uniqueness of the KAS-seq signal at the second CRISPR array and the absence of the same strong KAS peaks at the other two CRISPR arrays. The second CRISPR array is also separated from both the Cas6 gene and the KAS-seq peak by the s479 sRNA, the reported function of which is involved in the regulation of zinc transport ^55^; how this second array might fit into the overall picture is not clear. The functional significance of elevated chromatin accessibility over CRISPR arrays is also currently unknown. As prokaryotes exhibit an immense variety of CRISPR systems and number and organization of CRISPR arrays, mapping these properties in multiple other prokaryotes would be highly informative.

## Methods

Except where explicitly indicated otherwise, data was processed using custom-written Python scripts (https://github.com/georgimarinov/GeorgiScripts)

### *Haloferax volcanii* cell culture

*H. volcanii* cells were obtained from the DSMZ German Collection of Microorganisms and Cell Cultures GmbH (Cat # 3757), and cultured in *Halobacterium* media ^66,67^, prepared as follows: 7.50 g casamino acids, 10.00 g yeast extract, 3.00 g sodium citrate, 2.00 g KCl, 20.00 g MgSO_4_× 7 H_2_O, 0.05 g FeSO_4_ × 7 H_2_O, 0.20 mg MnSO_4_ × H_2_O, and 250.00 g NaCl were mixed with distilled water in a total volume of 1 L. Media was then autoclaved and allowed to cool.

*H. volcanii* was typically grown at 42 °C, except for where otherwise indicated.

Cultures were stored at room temperature when not actively growing.

### *Haloferax volcanii* genome assembly and annotations

For all analyses, the genome assembly and annotation for the *Haloferax volcanii* DS2 strain, downloaded from the NCBI database, and also matching the haloVolc1 version on the UCSC Microbial Genome Browser ^68^ (http:////microbes.ucsc.edu/), was used. The UCSC Microbial Genome Browser was used for visualization of genome browser tracks.

### ATAC-seq experiments

Several variations of the ATAC-seq assays were tested. As *H. volcanii* is an archaeon, i.e. a prokaryote without a nucleus, and as it does not have a cell wall (as many other prokaryotes do), the nuclei isolation step typical for ATAC-seq protocols used in eukaryotes was omitted.

For native ATAC-seq, cells (∼0.1, ∼1 or ∼10 × 10^6^ cells as measured by OD_600_) were pelleted at 10,000 *g* for 2 minutes, then resuspended in 50 *µ*L transposition mix (25 *µ*L 2× TD buffer, 2.5 *µ*L Tn5, 22.5 *µ*L ultrapure H_2_O), and incubated at 37 °C for 15 minutes. The reaction was stopped by adding 250 *µ*L PB Buffer (Qiagen, Cat # 28006) and purified using the MinElute PCR Purification Kit (Qiagen, Cat # 28006), eluting in 10 *µ*L EB buffer. PCR was carried out by mixing the 10 *µ*L eluate, 10 *µ*L H_2_O, 2.5 *µ*L i5 primer, 2.5 *µ*L i7 primer, and 25 *µ*L NEBNext High-Fidelity 2× PCR Master Mix, using the following thermocycler program: 3 minutes at 72 °C, 30 seconds at 98 °C, 10 cycles of: 98 °C for 10 seconds, 63 °C for 30 seconds, 72 °C for 30 seconds. Final libraries were purified using the MinElute PCR Purification Kit.

For crosslinked ATAC-seq, cells were fixed by adding 37% formaldehyde (Sigma) at a final concentration of either 0.1% or 1% and incubating for 15 minutes at room temperature. Formaldehyde was then quenched using 2.5 M glycine at a final concentration of 0.25 M. Cells were subsequently centrifuged at 10,000 *g* for 2 minutes, washed once in 1× PBS, and centrifuged again at 10,000 *g* for 2 minutes. Transposition was carried out as above for 15 minutes. The reaction was stopped with the addition of 150 *µ*L IP Elution Buffer (1% SDS, 0.1 M NaHCO_3_) and 2 *µ*L Proteinase K (Promega, Cat # MC5005), then incubated at 65 °C overnight to reverse crosslinks. DNA was isolated by adding an equal volume of 25:24:1 phenol:chloroform:isoamyl solution, vortexing and centrifuging for 3 minutes at 14,000 rpm, then purifying the top aqueous phase using the MinElute PCR Purification Kit, eluting in 10 *µ*L EB buffer. Libraries were generated as described above.

### ATAC-seq data processing

Demultipexed FASTQ files were mapped to the *H. volcanii* genome as 2 × 36mers using Bowtie ^69^ (version 1.0.1) with the following settings: `-v 2 -k 2 -m 1 --best --strata`. Duplicate reads were removed using `picard-tools` (version 1.99).

TSS scores were calculated as the ratio of ATAC signal in the region ±100 bp around TSSs versus the ATAC signal of the 100-bp regions centered at the two points ±2 kbp of the TSS as previously described ^70^.

Peak calling was carried out using MACS2^37^ with the following settings: `-g 4000000 -f BAM --to-large --keep-dup all --nomodel`.

### DNA isolation and naked DNA sequencing

Genomic DNA was isolated by centrifuging cells at 10,000 *g* and resuspending the pellet in 200 *µ*L 1 × PBS, then using the MagAttract HMW DNA Kit (Qiagen, Cat # 67563), following the manufacturer’s instructions.

Genomic DNA libraries were prepared using 5 ng of DNA in a 50-*µ*L transposition reaction (*x µ*L DNA, 22.5 - *x µ*L H_2_O, 25 *µ*L 2× TD buffer, 2.5 *µ*L Tn5). The reaction was carried out for 5 minutes at 55 °C, then stopped with 250 *µ*L PB buffer. DNA was isolated using the MinElute PCR Purification Kit and amplified as described above for ATAC-seq.

### NOMe-seq and dSMF experiments

NOMe-seq/dSMF experiments were carried out as previously described ^36^, with some modifications. Cells were pelleted at 10,000 *g*, then crosslinked as described for ATAC-seq at 1% formaldehyde concentration.

Fixed cells were resuspended in 100 *µ*L M.CviPI Reaction Buffer (50 mM Tris-HCl pH 8.5, 50 mM NaCl, 10 mM DTT), then treated with M.CviPI by adding 200 U of M.CviPI (NEB), SAM at 0.6 mM and sucrose at 300 mM, and incubating at 30 °C for 20 minutes. After this incubation, 128 pmol SAM and another 100 U of enzyme were added, and a further incubation at 30 °C for 20 minutes was carried out. For dSMF experiments, M.SssI treatment followed immediately, by adding 60 U of M.SssI (NEB), 128 pmol SAM, MgCl_2_ at 10 mM and incubation at 30 °C for 20 minutes. The reaction was stopped by adding an equal volume of Stop Buffer (20 mM Tris-HCl pH 8.5, 600 mM NaCl, 1% SDS, 10 mM EDTA).

Crosslinks were reversed overnight at 65 °C, and DNA was isolated using the MinElute PCR Purification Kit (Qiagen, Cat # 28006).

Enzymatically labeled DNA was then sheared on a Covaris E220, and converted into sequencing libraries following the EM-seq protocol, using the NEBNext Enzymatic Methyl-seq Kit (NEB, Cat # E7120L).

### NOMe-seq data processing

Adapters were trimmed from reads using Trimmomatic ^71^ (version 0.36). Trimmed reads were aligned against the *H. volcanii* genome using `bwa-meth` with default settings. Duplicate reads were removed using `picard-tools` (version 1.99). Methylation calls were extracted using `MethylDackel` (https://github.com/dpryan79/MethylDackel). Additional analyses were carried out using custom-written Python scripts (https://github.com/georgimarinov/GeorgiScripts).

### KAS-seq experiments

KAS-seq experiments were carried out following the previously published protocol ^32^ with some modifications in the sequencing library generation part.

Briefly, a 500-mM N_3_-kethoxal solution was brought to 37 °C, then added to 2 mL of culture at a final concentration of 5 *µ*M. Cells were then incubated for 5 minutes at 37 °C in a ThermoMixer at 1000 rpm.

Cells were then pelleted by centrifugation at 10,000 *g* for 1 minute, resuspended in 200 *µ*L 1 × PBS buffer, and DNA was immediately isolated using the Monarch Genomic DNA Purification Kit (NEB, Cat # T3010S), with the modification that elution was carried out with 50 *µ*L 25 mM K_3_BO_3_ solution (pH 7.0).

Biotin was clicked onto kethoxal-modified guanines by mixing 50 *µ*L DNA, 2.5 *µ*L 20 mM DBCO-PEG4-biotin (Sigma, Cat # 760749; DMSO solution), 10 *µ*L 10 × PBS, and 22.5 *µ*L 25 mM K_3_BO_3_ and incubating at 37 °C for 90 minutes.

DNA was isolated using AMPure XP beads and eluted in 130 *µ*L 25 mM K_3_BO_3_ (pH 7.0), then sheared on a Covaris E220 for 120 seconds down to ∼150-200 bp.

Libraries were built on beads using the NEBNext Ultra II DNA Library Prep kit (NEB, Cat # E7645L). Biotin pull down was initiated py pipetting 20 *µ*L Dynabeads MyOne Streptavidin T1 beads (ThermoFisher Scientific, Cat # 65306) into DNA lo-bind tubes. Beads were separated on magnet, resuspended in 200 *µ*L of 1 × TWB buffer (Tween Washing Buffer; 5 mM Tris-HCl pH 7.5; 0.5 mM EDTA; 1 M NaCl; 0.05% Tween 20), then separated on magnet again and resuspended in 300 *µ*L of 2 × BB (Binding Buffer; 10 mM Tris-HCl pH 7.5, 1 mM EDTA; 2 M NaCl). The DNA (130 *µ*L) was added together with 170 *µ*L 0.1 × TE buffer, and incubated at RT on rotator for ≥ 15 minutes. Beads were separated on magnet, resuspended in 200 *µ*L of 1× TWB, and incubated at 55 °C in a Termomixer for 2 minutes with shaking at 1000 rpm. Beads were again separated on magnet and the 200-*µ*L 55 °C TWB wash step was repeated. Beads were separated on magnet and resuspended in 50 *µ*L 0.1× TE.

End repair was carried out by adding 7 *µ*L NEB End Repair Buffer and 3 *µ*L NEB End Repair Enzyme, incubating at 20 °C for 30 minutes, then at 65 °C for 30 minutes.

End repair was followed by adaptor ligation by adding 2.5 *µ*L NEB Adaptor, 1 *µ*L NEB Ligation Enhancer and 30 *µ*L NEB Ligation Mix, incubating at 20 °C for 20 minutes, then adding 3 *µ*L USER Enzyme and incubating at 37 °C for 15 minutes. Beads were separated on magnet, resuspended in 200 *µ*L of 1 × TWB, then incubated at 55 °C in a Thermomixer for 2 minutes with shaking at 1000 rpm. Subsequently beads were separated on magnet and resuspended in 100 *µ*L of 0.1× TE, separated on magnet again, resuspended in 15 *µ*L of 0.1 × TE Buffer, and transfered to PCR tubes.

Beads were then incubated at 98 °C for 10 minutes, and libraries were amplified by adding 5 *µ*L of i5 primer, 5 *µ*L of i7 primer and 25 *µ*L of 2× Q5 Hot Start Polymerase Mix, using the following PCR program: 30 seconds at 98 °C; 15 cycles of 98 °C for 10 seconds, 65 °C for 30 seconds, and 72 °C for 30 seconds; and a final extension at 72 °C for 5 minutes.

Beads were separated on magnet and the final libraries were purified from the supernatant using 50 *µ*L AMPure XP beads, eluting in 0.1× TE buffer.

### KAS-seq data processing

Demultipexed FASTQ files were mapped to the *H. volcanii* genome as 2 × 36mers using Bowtie ^69^ with the following settings: `-v 2 -k 2 -m 1 --best --strata`. Duplicate reads were removed using `picard-tools` (version 1.99).

### Multimapping reads analysis

For the purpose of examining repetitive regions in the genome, such as the rRNA operons, which exist in two identical copies in the genome, and are thus not uniquely mappable, reads were mapped with the `-a` option instead of `-k 2 -m 1`. Normalization was carried out as previously described and discussed ^72^.

### Differential accessibility/KAS-seq analysis

The analysis of differential chromatin accessibility as measured using ATAC-seq or enriched for KAS-seq signal was carried out using DESeq2^73^. Read counts were calculated over promoters or gene bodies and used as input into DESeq2.

### External sequencing datasets

MNAse-seq datasets for *H. volcanii* were downloaded from NCBI accession PRJNA174818^22^, and processed as described above for ATAC-seq and KAS-seq.

ATAC-seq for *Suflolobus islandicus* ^40^ was downloaded through the Short Read Archive (SRA) from BioProject accession 814106.

## Data availability

The sequencing datasets generated for and used in this study can be accessed from GEO accession GSE207470.

## Author contributions

G.K.M. conceptualized the study, carried out cell culture, performed ATAC-seq, KAS-seq, and NOMe-seq/dSMF, analyzed the data and wrote the manuscript, with input from W.J.G. S.T.B. performed ATAC-seq, KAS-seq, and NOMe-seq/dSMF experiments. T.W. and C.H. provided key reagents. A.K. and W.J.G. supervised the study.

## Acknowledgments

The authors would like to thank Samuel Kim for input on the KAS-seq protocol, Matthew P. Swaffer for technical assistance, Zohar Shipony and members of the Greenleaf and Kundaje labs, and Tobias Warnecke, Fabian Blombach and Anita Marchfelder for helpful discussions and suggestions for improvements to the text and analysis. This work was supported by NIH grants (P50HG007735, RO1 HG008140, U19AI057266 and UM1HG009442 to W.J.G., 1UM1HG009436 to W.J.G. and A.K., 1DP2OD022870-01 and 1U01HG009431 to A.K.), the Rita Allen Foundation (to W.J.G.), the Baxter Foundation Faculty Scholar Grant, and the Human Frontiers Science Program grant RGY006S (to W.J.G). W.J.G is a Chan Zuckerberg Biohub investigator and acknowledges grants 2017-174468 and 2018-182817 from the Chan Zuckerberg Initiative.

## Competing interests

W.J.G. is a consultant and equity holder for 10x Genomics, Guardant Health, Quantapore, and Ultima Genomics, and cofounder of Protillion Biosciences, and is named on patents describing ATAC-seq.

A.K. is a consulting Fellow with Illumina, a member of the SAB of OpenTargets (GSK), PatchBio, SerImmune and a scientific co-founder of RavelBio.

## Supplementary Materials

### Supplementary Figures

**Supplementary Figure 1:**
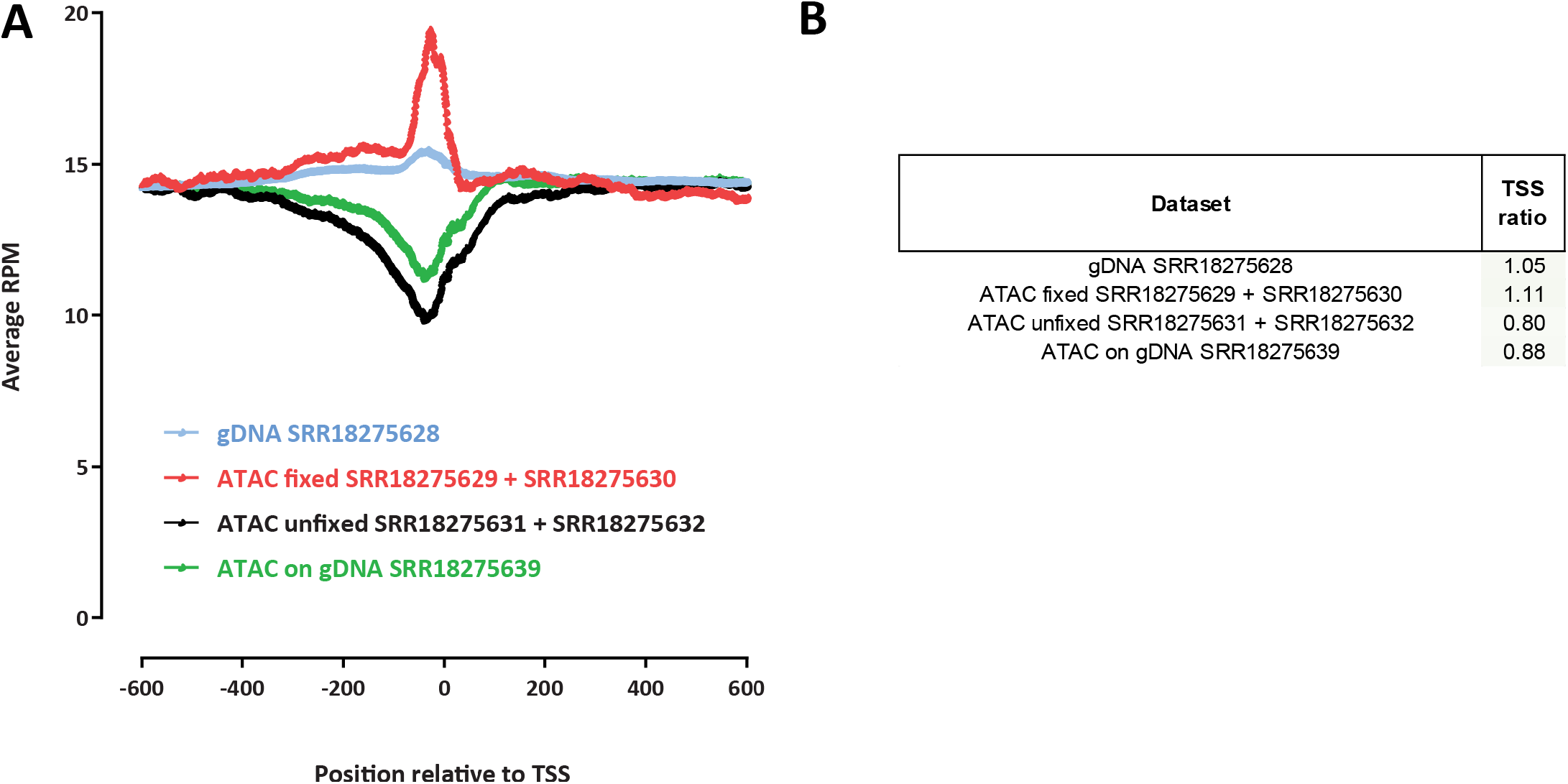
TSS enrichment levels in ATAC-seq data for *Suflolobus islandicus*. (A) TSS metaprofiles for ATAC-seq in fixed and unfixed cells and for genomic DNA libraries. (B) TSS scores for each dataset.

**Supplementary Figure 2:**
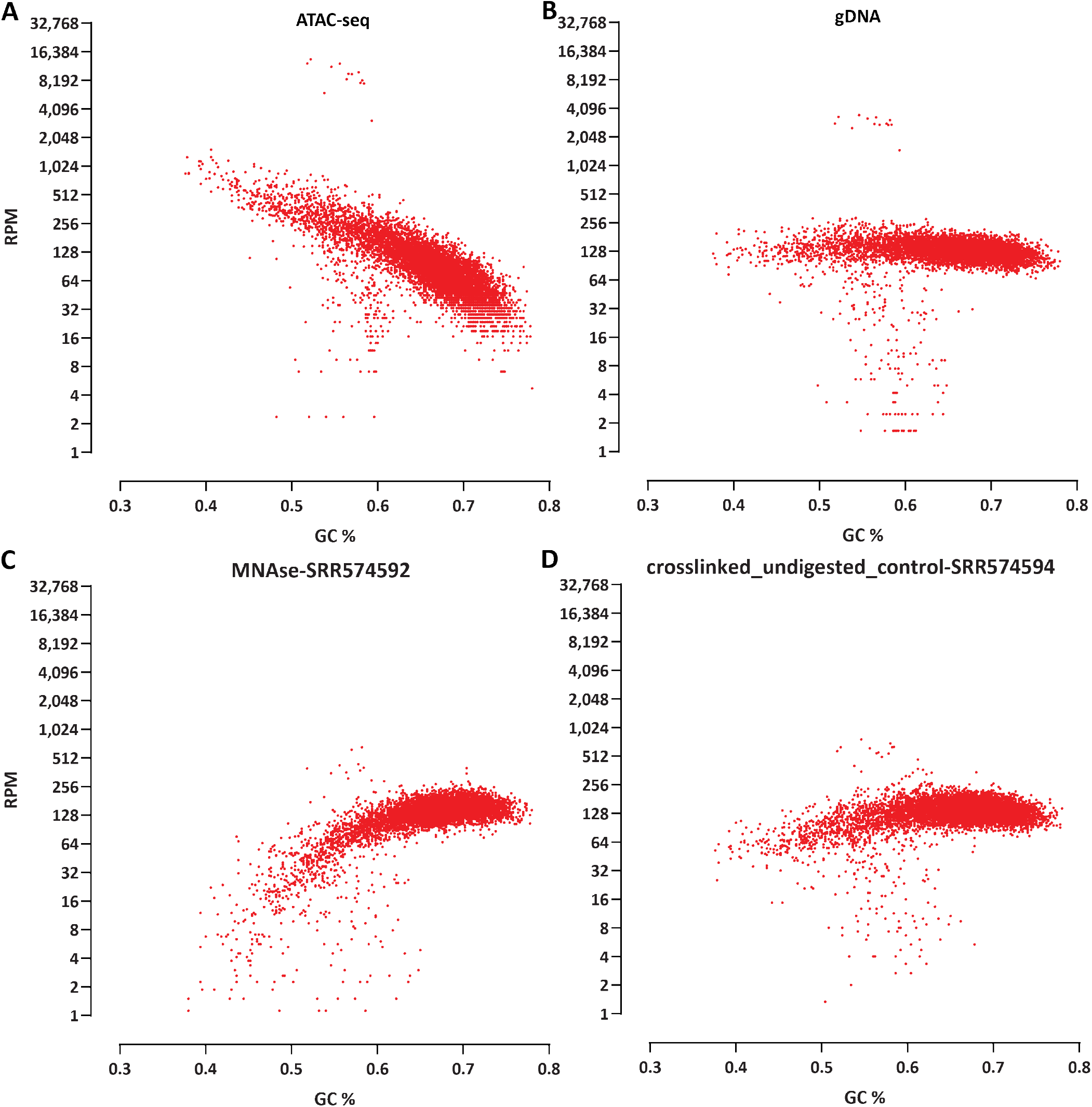
Correlation between the extent of chromatinization and GC content in *Haloferax*. The genome was split in 500-bp bins and RPM values and GC% were calculated for each bin. (A) ATAC-seq; (B) Tagmented naked genomic DNA control; (C) MNase-seq (external dataset); (B) Crosslinked undigested control (external dataset).

**Supplementary Figure 3:**
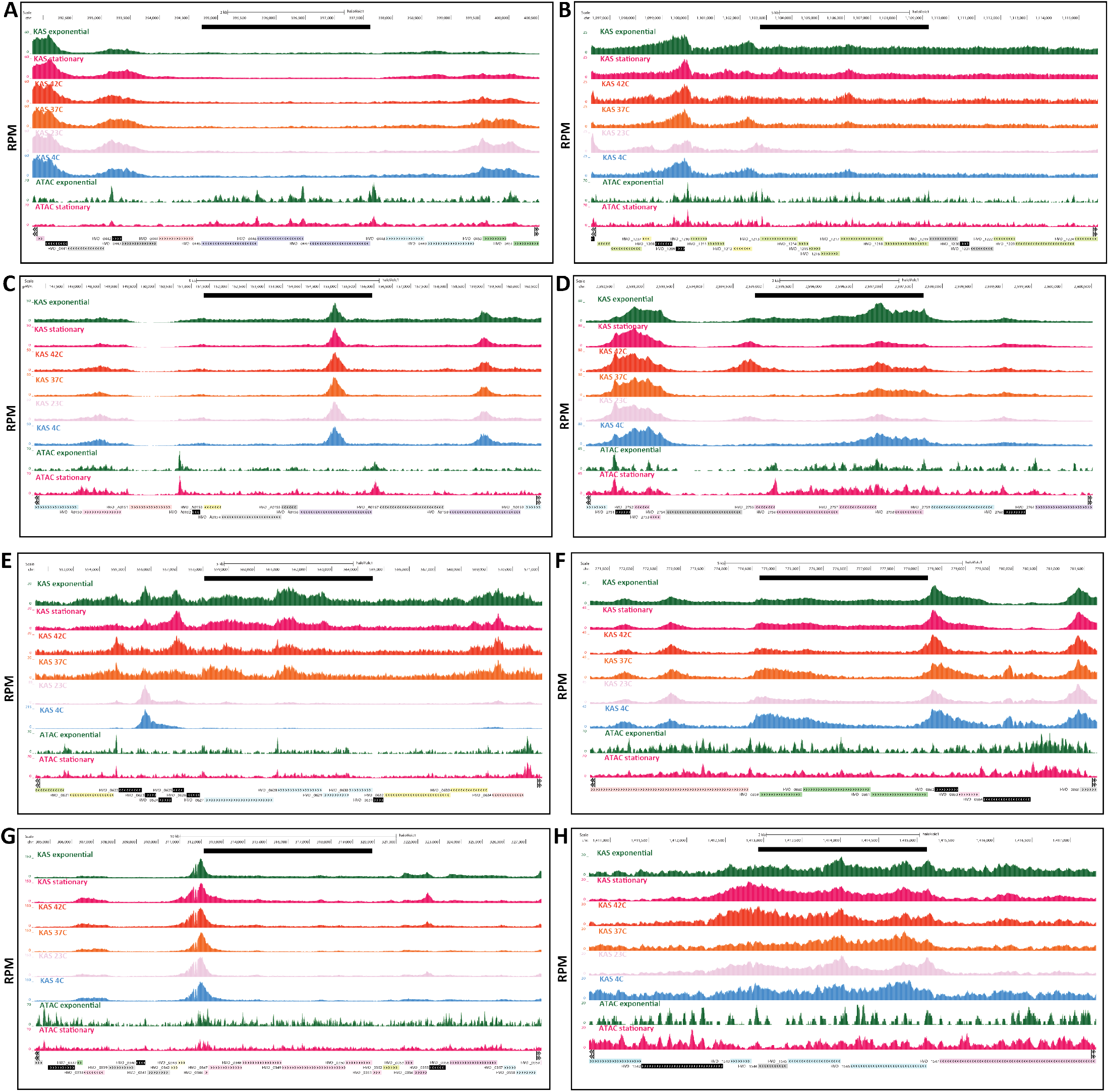
Coordination between chromatin accessibility and transcriptional activity within *H. volcanii* operons. Black bar shows the operon boundaries. (A) Putative phosphate-phosphonate ABC transporter. (B) Flagellar cluster (FlaCE, FlaF, FlaG, FlaH, FlaI, FlaJ). (C) Urease accessory protein operon (UreG, UreD, UreE, UreF). (D) 50S ribosomal proteins L12, L10, L1, L11. (E) Putative ABC transporter. (F) FeS assembly genes SufC, SufB, SufD. (G) RNA Polymerase subunits. (H) Dihydroxyacetone kinase subunit L, subunit DhaK, and phosphotransfer subunit.

**Supplementary Figure 4:**
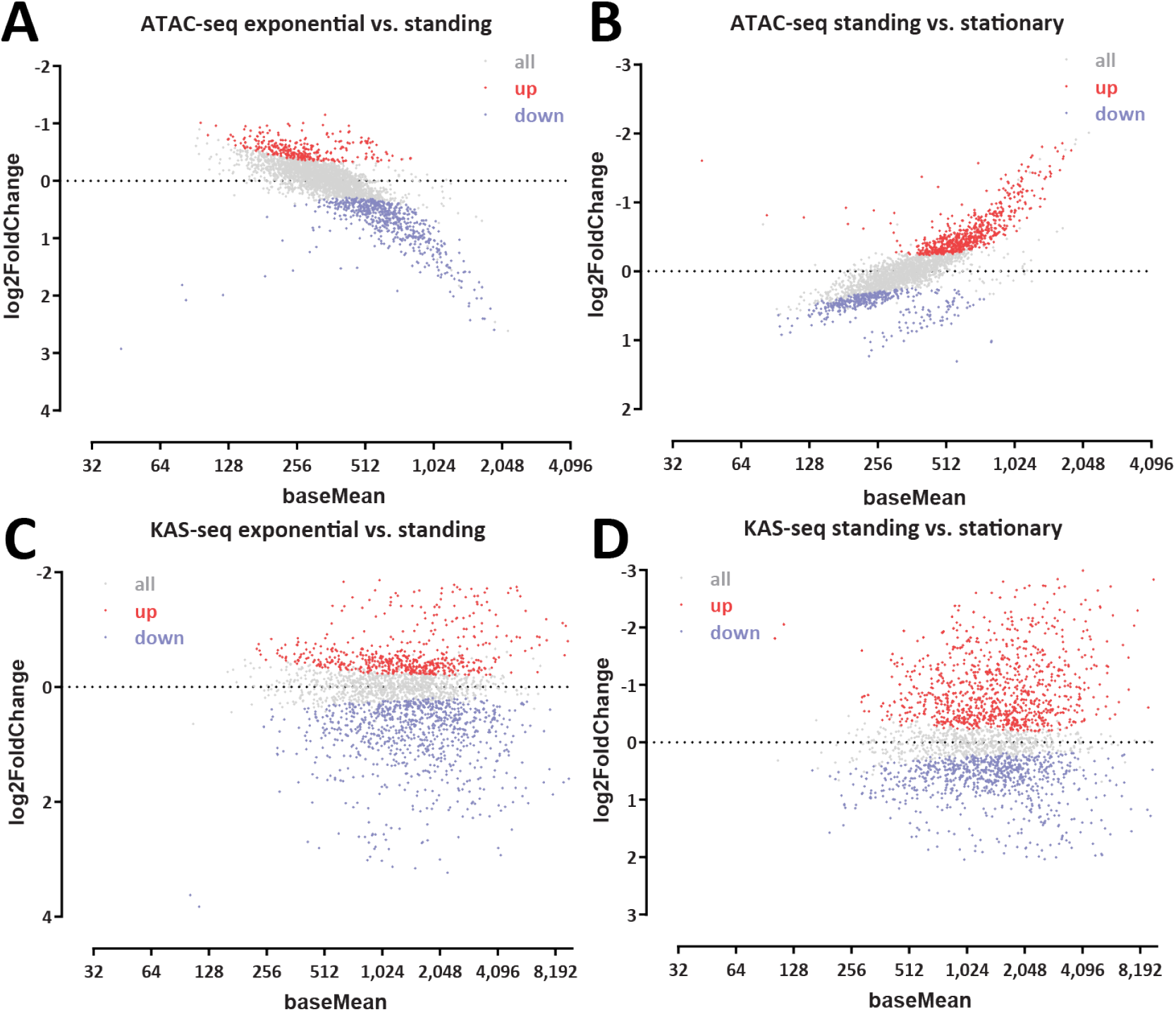
Differential ATAC-seq in KAS-seq analysis for standing *H. volcanii* cells. (A-B) Differential chromatin accessibility; (C-D) Differential KAS-seq levels.

## Notes

### Summary of Updates

The version has been updated to include some new results and a revised interpretation of certain findings in it.

